# Detection of Fortunate Molecules Induce PArticle Resolution Shift (PAR-shift) towards Single-molecule Limit in SMLM: A Technique for Resolving Molecular Clusters in Cellular System

**DOI:** 10.1101/2022.03.22.485352

**Authors:** S Aravinth, Prakash Joshi, Partha P. Mondal

## Abstract

Molecules capable of emitting a large number of photons (also known as fortunate molecules) are crucial for achieving resolution close to a single molecule limit (the actual size of a single molecule). We propose a long-exposure single molecule localization microscopy (leSMLM) technique that enables detection of fortunate molecules, which is based on the fact that detecting a relatively small subset of molecules with large photon emission increases its localization 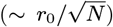. Fortunate molecules have the ability to emit a large burst of photons over a prolonged time (*>* average triplet-state lifetime). So, a long exposure time allows the time window necessary to detect these elite molecules. The technique involves the detection of fortunate molecules to generate enough statistics for a quality reconstruction of the target protein distribution in a cellular system. Studies show a significant PArticle Resolution Shift (PAR-shift) of about 6 *nm* and 11 *nm* towards Single-molecule-limit (away from diffraction-limit) for an exposure time window of 60 *ms* and 90 *ms*, respectively. In addition, a significant decrease in the fraction of fortunate molecules (single molecules with small localization precision) is observed. Specifically, 8.33% and 3.43% molecules are found to emit in 30 − 60 *ms* and 60 − 90 *ms*, respectively, when compared to SMLM. The long exposure has enabled better visualization of Dendra2HA molecular cluster, with sub-clusters within a large cluster. Thus, the proposed technique (*leSMLM*) facilitates a better study of cluster formation in fixed samples. Overall, the method enables better spatial resolution at the cost of relatively poor temporal resolution.

---

Fortunate molecules can transform single-molecule biology and is a potential candidate for pushing resolution close to single-molecule limit (the actual size of single-molecule) [1]. In general, SMLM typically provides resolution somewhere in the spatial range of 30 − 50 *nm*, but seldom do they provide true molecular resolution [14–19]. Recent advances in super-resolution microscopy (e.g., STED, MINFLUX, dSTORM, SIMPLE and ROSE) have shown sub-10 nm resolution [2] [3] [4] [5] [6] [8]. For achieving high resolution in single-molecule localization microscopy (SMLM), molecules with large photon emission (fortunate molecules) are likely to play a crucial role [1]. This requires selective detection of very bright molecules (decided by the product of fluorescence quantum yield and the molar extinction coefficient) for labelling (e.g., pamCherry, Dendra2, Cy5, and Alexa Fluor 647) the protein of interest [9] [7]. Another possibility is to increase the detection time window of the detector to capture a small set of elusive fortunate molecules. Nonetheless, a plethora of prospects are likely to emerge with the capability of improved localization precision of single molecules. This will push the particle resolution (PAR) towards the single-molecule limit.

Since its first inception, the field of super-resolution imaging has proliferated, and the technique is finding new applications in the broad field of bioimaging and biophysics [13–20]. The last decade has seen many variants including, ground-state depletion microscopy (GSDIM) [21], super-resolution optical fluctuation imaging (SOFI) [22], points accumulation for imaging in nanoscale topography (PAINT) [23] [24], simultaneous multiplane imaging-based localization encoded (SMILE) [25] [26], individual molecule localization–selective plane illumination microscopy (IML-SPIM) [27], MINFLUX [8],POSSIBLE microscopy [1] and others [28–35]. The reconstructed super-resolution image forms the basis for understanding biophysical mechanisms, 3D organization of organelles, multiprotein complexes, and dynamical events in cellular systems. Therefore, methods that improve resolution within the existing hardware of optical imaging configurations are in high demand. Such techniques are likely to be accepted by researchers across the science spectrum (from Biology to physics).

Existing SMLM techniques involve recording several frames at video rate (approximately 16 frames/sec). This translates into 30 milliseconds exposure time per frame which coincides roughly with the triplet state lifetime of target single molecules [15]. Although this is an average triplet state lifetime, the rates differ depending on the local chemical environment and/or the presence of other heavy metals. For example, it is noted that the lifetime of fluorescent probes becomes longer as pH is shifted to the acidic environment [36]. In another study, fast quenching of the fluorescence for Alexa647 is observed due to its interaction with aromatic amino acid residues [37]. Considering these factors and similar observations [15], we wish to investigate molecules with large excitation-emission cycles, i.e., those that emit a relatively large number of photons. One way to collect these elusive molecules is to increase the exposure time of the detection process. This allows a larger time window to detect all the photons emitted by these elusive molecules. So, the strategy is to look for fortunate molecules, that have better localization precision given by, 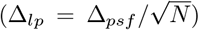, where, Δ_*psf*_ and *N* are the diffraction-limited PSF and number of detected photons, respectively [38]. The technique is expected to improve the overall resolution of the reconstructed image, which directly affects the resolvability of large molecular clusters formed in the complex cellular system.

This letter proposes a method to better the spatial resolution of localization-based super-resolution microscopy by sacrificing the temporal resolution. Thus our techniques are better suited for fixed samples rather than live specimens. Subsequently, the method is explored that showed resolvability of large molecular clusters indicating the presence of sub-clusters. These parameters are then used for determining critical biophysical parameters (cluster density, area, and the number of molecules) in a cellular system.

We study the influenza type-A model that involves dynamic clustering of the glycoprotein Hemagglutinin (HA) in transfected NIH3T3 cells. HA protein is an antigenic glycoprotein found on the surface of influenza viruses and is responsible for binding the virus to the cell. Cells accumulate these proteins based on the local physiology during the onset of viral infection (Influenza A) [39] [40] [41]. It may be noted that HA clustering is a key process and have a direct bearing on the infectivity rate. So, it becomes essential to understand the HA accumulation process and find methods to disrupt it. In the present study, the cells were transfected with Dendra2HA plasmid DNA and cultured for 24 hours before fixing them for superresolution imaging.

The Dendra2 molecule involves the conversion between two states (on-off) mediated by triplet state [10] [11]. Accordingly, the priming or excitation of the anionic cis-chromophore populates the S1-state [42] [12]. The de-population occurs via either fluorescence pathway or low-yield intersystem crossing to the lowest triplet state, T1. The mechanism involving conversion of primed state strongly relies on the creation of a triplet intermediate state. This results in fluorescence intermittencies such as triplet state transitions (popularly known as blinking) that are observed in many molecules. In general, several photoactivable fluorescent proteins show blinking timescale of a few tens of milliseconds (e.g., Alexa555 has a timescale of 22 ± 6 *ms* in deoxygenated PBS buffer [43], and Dendra2 has an average bleaching rate of 23.1 ± 1.9 *s*^−1^ [44]). This necessitates that the images be acquired between 15 − 20 frames / sec. While a significant fraction of photoswitches is non-fluorescent, a small subset is stochastically activated and localized. The molecules can be localized to high precision by fitting a Gaussian to their point spread function and determining the number of collected photons. This leads to high precision localization of single molecules to render a super-resolution map of the specimen. While molecules are known to blink in the time window 20 − 35 *ms*, a good fraction of molecules blink for longer timescales [15]. So, blinking that occurs at large timescales (*>* 35 *ms*) is averaged out in the process. Here, we observed blinking for relatively longer timescales (up to 90 *ms*). This allows for the detection of a large number of molecules with a longer fluorescence cycle that enables better localization precision. Of course, this allows more time that may cause significant movement of the molecules in a dynamic cellular environment. Hence, long exposure-based techniques appear to be better suited for specimens where the timescale for molecular dynamics is relatively larger than (*τ*_*exp*_).

The schematic diagram of the *leSMLM* optical system and major components is shown in Fig. 1. The technique employs two lasers: 405 nm laser for activation and 561 nm laser for excitation of the target photoactivable molecule (Dendra2-HA). The beams are combined using a dichroic mirror (DMLP425R, Thorlabs, USA) and directed to the high NA objective lens. In addition, a third optical subsystem (widefield optical setup) is also coupled to the leSMLM system for identifying transfected cells. It involves blue light (*λ* = 470 − 490 *nm*) illumination for identifying transfected cells (transfected with Dendra2-HA protein), which predominantly emits in the Stoke-shifted wavelength 517 *nm*. Superresolution imaging is achieved by subjecting the cell to 405 nm and 561 nm light for activation and excitation of Dendra2-HA. An objective lens of high numerical aperture (*Olympus*, 1.3*NA*, 100*X*) is used to illuminate the specimen, and the same is used to collect the fluorescence at *λ >* 575 *nm*. The fluorescence is then directed to the detection subsystem that magnifies the image by 266*X* and filters the fluorescence light. The filter box consists of two notch filters (NF03-405E-25 and NF03-561E-25, Semrock, USA) and a long-pass filter (FF01-593/46-25, Semrock, USA) to filter-out illumination light and background radiation. To detect single molecules, exposure times of 30 *ms*, 60 *ms*, and 90 *ms* are considered, and 3871 − 8085 images are recorded. Subsequently, the images are analyzed (background subtraction, spot detection, and Gaussian fit to the single-molecule PSF), and parameters (localization precision, the number of photons, among others) are estimated. All these parameters are then used to identify clusters and determine biophysical parameters.

**FIG. 1:**
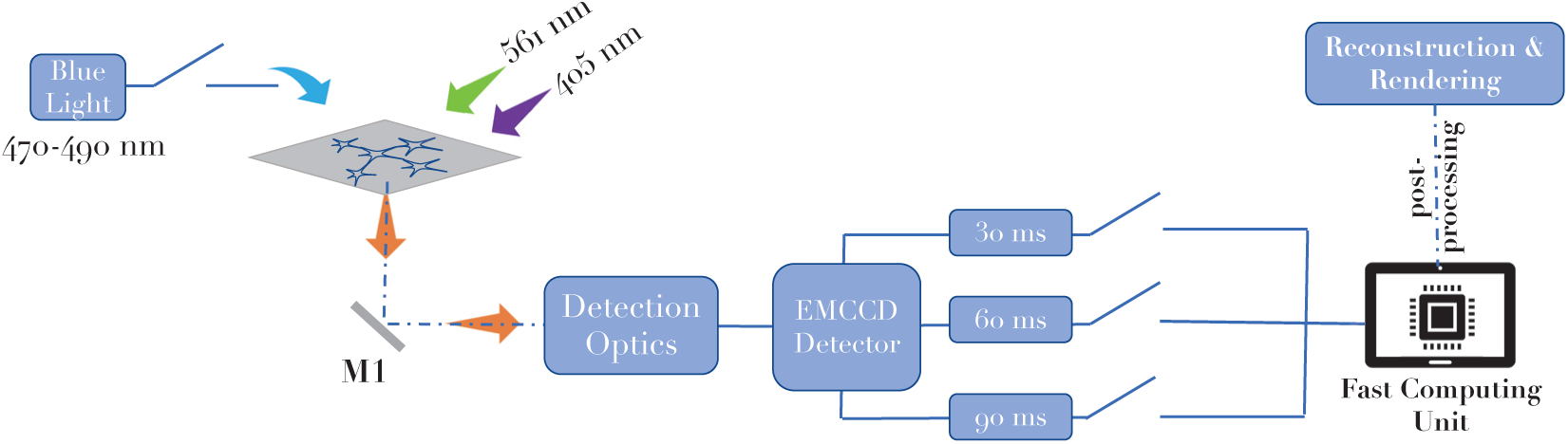
Schematic diagram for the proposed *leSMLM* super-resolution scheme to detect fortunate molecules. Blue light is used to identify transfected cells, whereas the combination of 405 *nm* and 561 *nm* is used for activating and exciting single molecules. The molecules are recorded by EMCCD detector at 30, 60, 90 *ms* exposure and subsequently processed by a fast computing unit.

Fig. 2 shows the transfected cells among other non-transfected ones. We observed a transfection efficiency of about 14% at 24 *hrs*. The corresponding super-resolved image reconstructed from 4085 images with a total of 19132 single-molecule signatures is shown along with the localization precision. The map shows the existence of molecular clusters of Dendra2-HA molecules. Both standard SMLM (*τ*_*exp*_ = 30 *ms*) and leSMLM (*τ*_*exp*_ = 60 *ms*) are used to understand the nature of these clusters (see, Fig. 3). Specifically, one can notice the resolvability of HA clusters for leSMLM compared to standard SMLM. This is because fortunate molecules allow better localization when compared to single molecules. Large exposure time-window (at 60 *ms*) has facilitated the detection of these elusive molecules that are capable of emitting large number of photons, thereby significantly improving the localization precision 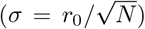, and thereby reducing the effective size of the Gaussian representing single molecules. The observation is significant since this demonstrates the ability of *leSMLM* to identify sub-molecular clusters formed during the early stages of infection. This is better illustrated by the localization precision plots in Fig.4, which shows a significant shift in Particle Resolution (termed as, PAR-shift). Specifically, we observed a shift of 6 *nm* and 11 *nm* towards the single-molecule-limit for leSMLM (with an exposure time of 60 *ms* and 90 *ms*) when compared to standard SMLM. To better understand PAR-shift observed at large exposure time, we analyzed localization precision in parts. Fig. 5 shows the localization plots for six chosen size regimes, each 10 *nm* apart. In the sub-10 *nm* and 10 − 20 *nm* regime, there is a predominant presence of fortunate molecules (see, Green bars in Fig. 5 A-C), while the number falls sharply for *>* 30 *nm* (see, red bars in Fig. 5 D-F). This can be also be inferred from the %overlap in the localization plots. Here, overlap is defined as the ratio of # molecules having the same localization precision for SMLM and leSMLM by the total # of molecules. This simply indicates the dominance of fortunate molecules (indicated by green bars in Fig. 5 A-C) for *<* 30 *nm* range. So, the detection of fortunate molecules causes PAR-shift towards the actual single-molecule limit (*a*_0_).

**FIG. 2:**
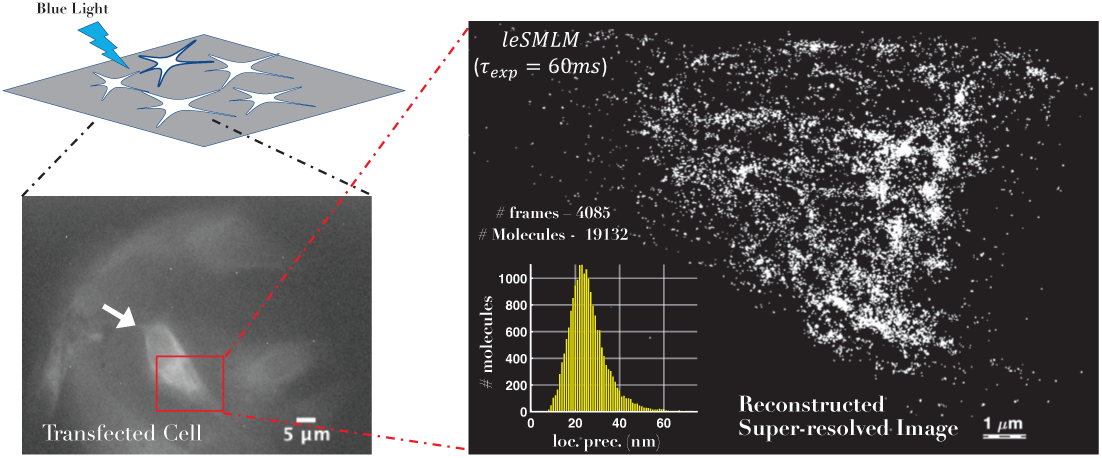
*leSMLM* super-resolved image of a Dendra2HA transfected NIH3T3 cell. Alongside, transfected cell image and localization precision are also shown.

**FIG. 3:**
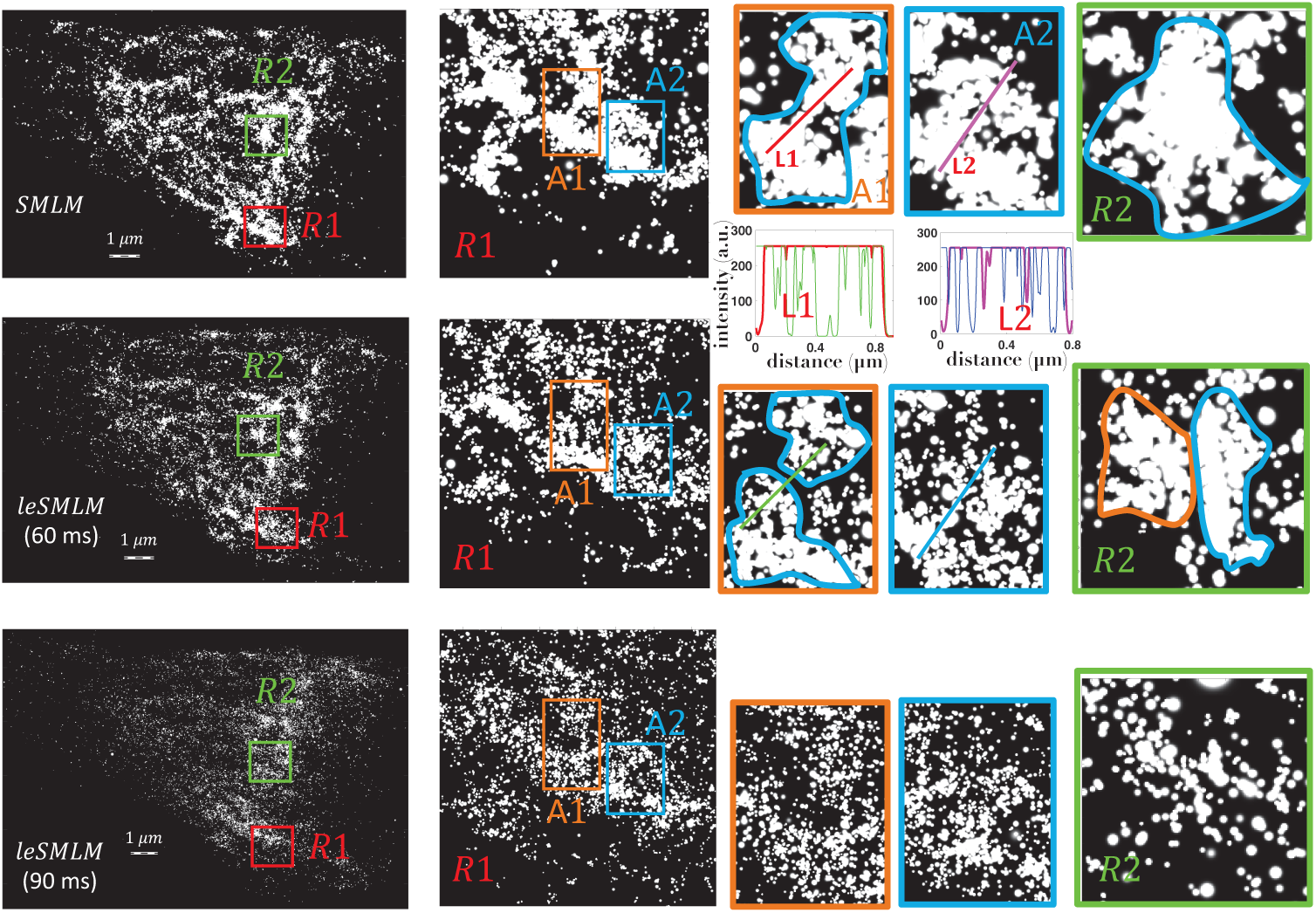
Reconstructed super-resolved image for leSMLM (*τ*_*exp*_ = 60 *ms* and *τ*_*exp*_ = 90 *ms*). Alongside HA clusters (marked by color contours) are identified and intensity plots (L1 and L2) are shown. For comparison, SMLM reconstruction (*τ*_*exp*_ = 30 *ms*) is also shown. Resolve molecular clusters in leSMLM (*τ*_*exp*_ = 60 *ms*) is quite evident.

**FIG. 4:**
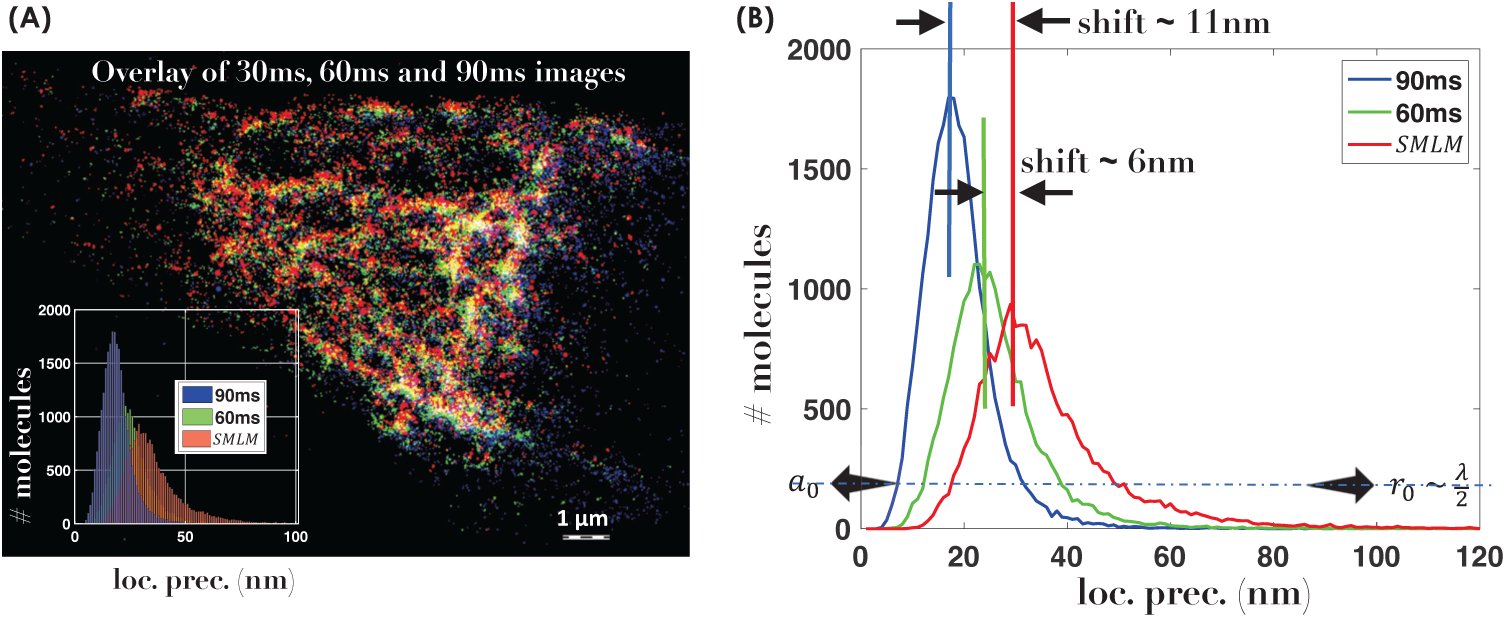
Overlay of SMLM and leSMLM images (*τ*_*exp*_ = 60 *ms* and *τ*_*exp*_ = 90 *ms*), displaying HA clusters at varying resolution. Localization precision plots show a PAR-shift of 6 *nm* and 11 *nm* towards single-molecule-limit (*a*_0_), and away from diffraction limit (*r*_0_ *∼ λ/*2).

**FIG. 5:**
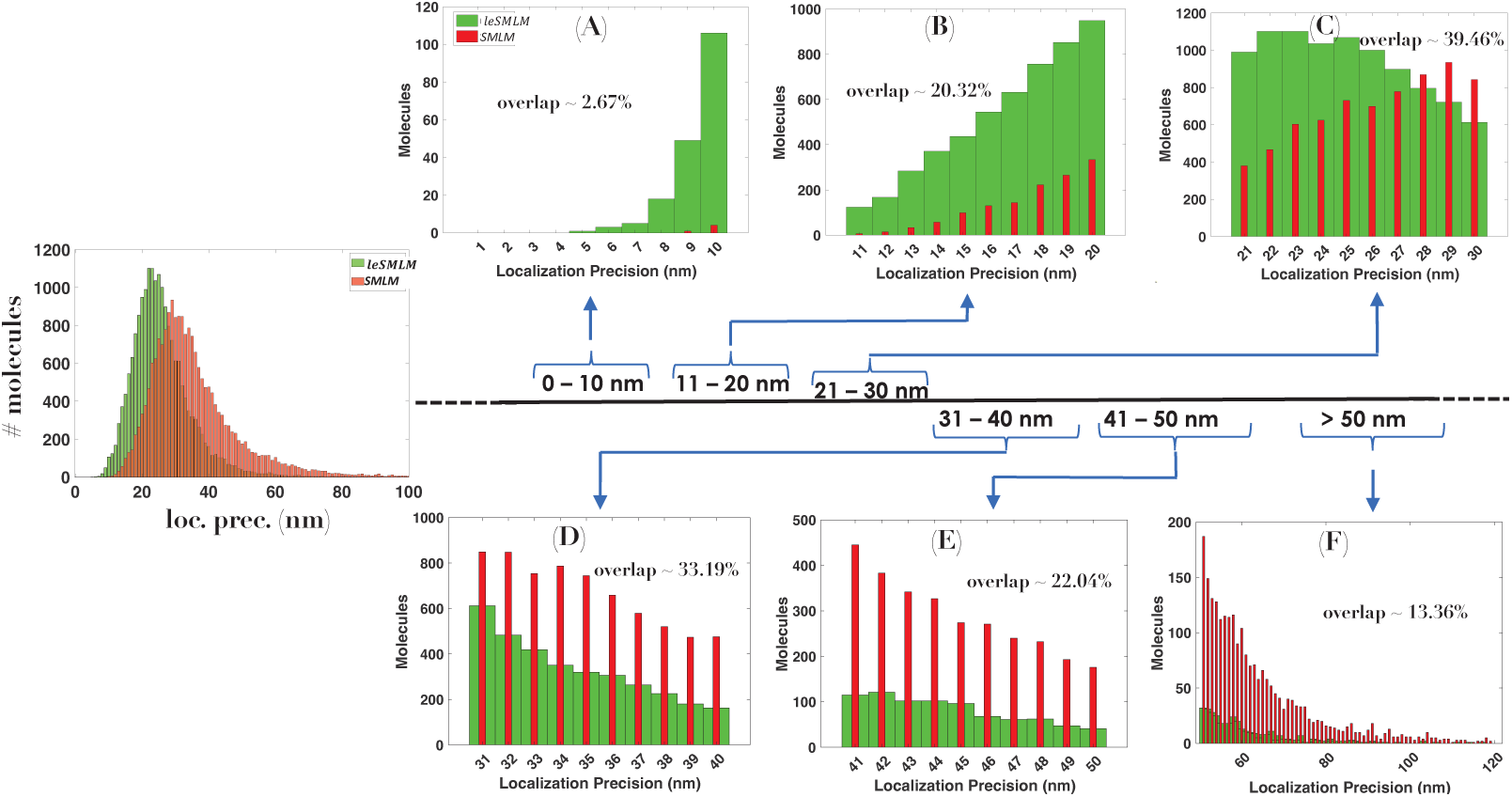
Region-wise segregation of localization data showing a significant increase in fortunate molecules specifically in the 0 − 10 *nm*, 10 − 20 *nm* and 20 − 30 *nm* regime. Corresponding percentage overlap are also shown.

Next, we concentrate on the Dendra2-HA cluster formation in the transfected NIH3T3 cells to understand the biological significance of HA dynamics. Fig. 6 shows cluster analysis on leSMLM and standard SMLM reconstructed super-resolved images. It is immediately evident that the number of clusters identified by the point clustering algorithm has increased to 33 compared to 27 for standard SMLM. This is because leSMLM can better resolve large clusters, indicating the presence of sub-clusters. We further carried out parameter estimation and determined critical biophysical parameters such as cluster density, size (area), and the number of molecules per cluster. The corresponding histogram plots for these parameters are shown in Fig. 6 (insets). General parameters characterizing HA cluster are tabulated in Table. 1. It may be noted that the average number of molecules per cluster and its size has decreased while the density of clusters has remained similar. This indicates an increase in the number of clusters that include sub-clusters (a total of 33 for leSMLM) compared to relatively fewer clusters (about 27) obtained for standard SMLM. However, the downside of leSMLM is the poor temporal resolution that calls for a relatively large number of frames (equivalently, more molecules) for quality reconstruction.

**TABLE I:** Key biophysical parameters such as, mean # mol. /cluster (M/C), mean density (D), mean cluster area (A).

| $\tau_{exp}$ | #frames | #molecules | #clusters | M/C | D | A |
| --- | --- | --- | --- | --- | --- | --- |
| (ms) | . | . | . | . | #/$\mu m^2$ | $\mu m^2$ |
| 30 | 3871 | 20184 | 27 | 658 | 551 | 1.54 |
| 60 | 4085 | 19132 | 33 | 505 | 576 | 1.05 |
| 90 | 8085 | 25012 | - | - | - | - |

**FIG. 6:**
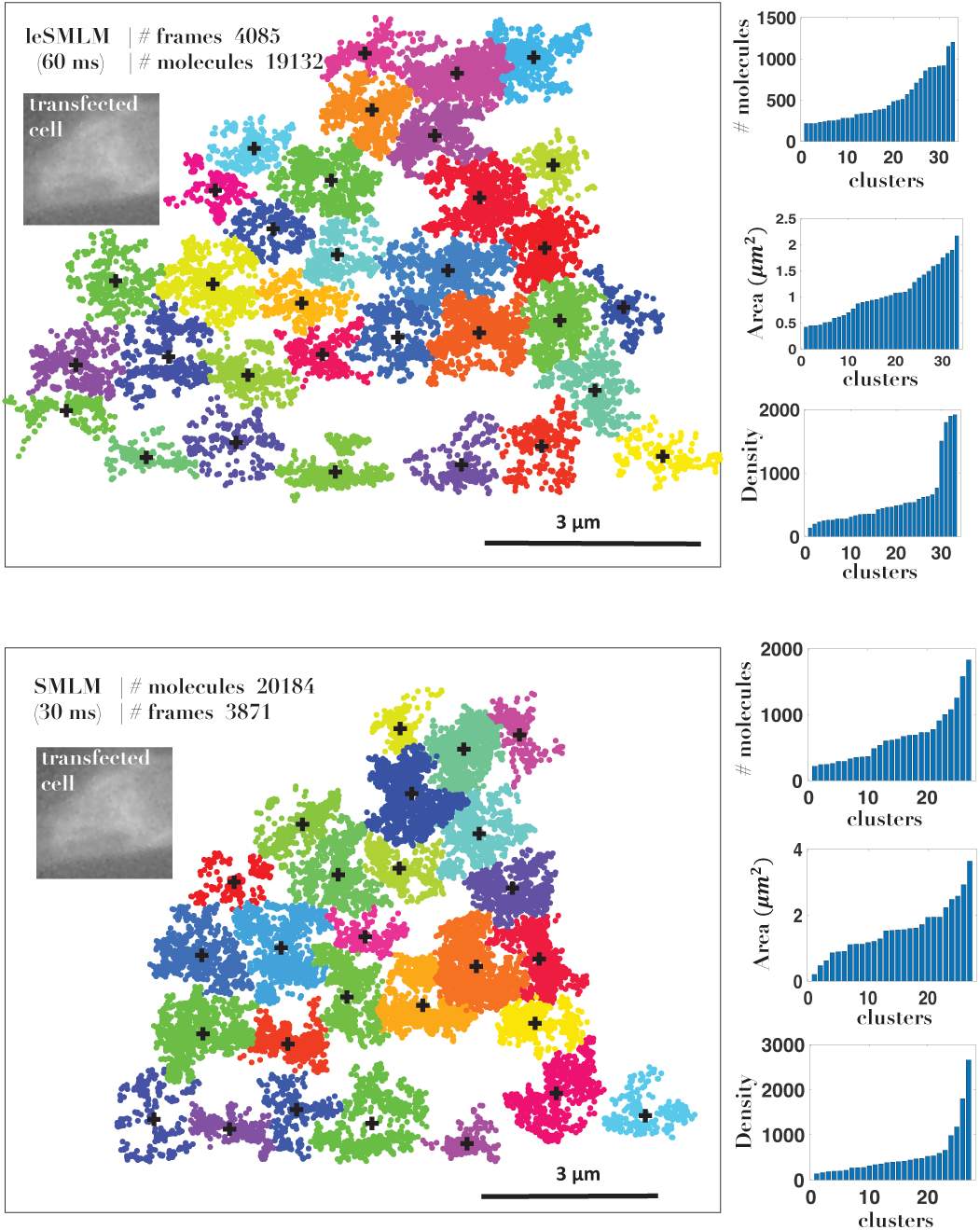
Point clustering analysis for traditional SMLM and leSMLM microscopy. Alongside, biophysical parameters related to the HA cluster (density, # mol./cluster, area) are also displayed.

We demonstrated an optical imaging technique (leSMLM) for better resolvability of HA clusters in Dendra2-HA transfected NIH3T3 cells that revealed the presence of sub-clusters. This is made possible by the detection of fortunate molecules with better localization precision, and as a result, the Gaussian-fit representing single molecules has a small size. Compared to the standard SMLM, the proposed technique increased the fraction of fortunate molecules, specifically in the sub-10 nm regime. This has helped distinguish sub-clusters hidden within large molecular clusters, which appear as a single large cluster. Such a process has biological significance, especially in disease biology, where the formation of small clusters leads to large clusters of viral HA protein in the infected cell (here, NIH3T3 cells). Moreover, standard classification algorithms can be used for determining essential characteristics of HA clusters related to their size, density, and the number of molecules. However, the leSMLM requires more data than SMLM to provide valuable information. So the system suffers from relatively poor temporal resolution, making it better suited for a fixed sample. Overall, it is evident that leSMLM can resolve features better than SMLM with the added benefits of quality cluster analysis.

## Acknowledgements

Financial funding from parent institute (Indian Institute of Science) is highly acknowledged.

## Data Availability

The data that support the findings of this study are available from the corresponding author upon request.

